# Maternal *Trypanosoma cruzi* infection is associated with significant placental remodeling regardless of vertical transmission

**DOI:** 10.64898/2026.01.13.699142

**Authors:** Sneider Alexander Gutierrez Guarnizo, Jaime So, Carolina Duque, Paloma Samame, Jean Karla Velarde, Emily Arteaga, Clariza Roxana, Luciana Basma, Liseth Roque, Jessy Condori, Martín Obregón, Edith Malaga, Jill Hakim, Shenice Harrison, Rony Colanzi, Manuela Verastegui, Monica Pajuelo, Freddy Tinajeros, Robert Gilman, Natalie Bowman, Monica R. Mugnier, The Chagas Disease Working Group in Bolivia and Peru

## Abstract

Chagas disease is a major protozoan infection in the Americas, causing approximately 12,000 deaths each year. It is caused by *Trypanosoma cruzi*, and can be transmitted transplacentally, leading to congenital Chagas disease, a silent route that carries substantial risk for newborns. However, the mechanisms underlying congenital Chagas transmission are poorly understood. Here, we evaluated whether *T. cruzi* infection alters the placental microenvironment and systemic physiology, and whether such alterations are associated with congenital transmission. Integrating bulk RNA sequencing, proteomics, and spatial transcriptomics, we show that *T. cruzi* infection elicits profound molecular remodeling in both placenta and peripheral blood, regardless of transmission status. Transmitting mothers exhibit a distinct transcriptional signature enriched for inflammatory and tissue-remodeling pathways. Notably, peripheral blood profiles mirrored some placental alterations. A panel of inflammatory serum proteins showed promising predictive potential for transmission risk, with implications for prenatal monitoring. Together, these findings support a fundamental shift in the conceptual framework of congenital Chagas disease, from a transmission-centered model to one that recognizes infection-driven placental damage as a pathological spectrum and identifies peripheral blood as a promising, non-invasive source of predictive biomarkers for adverse pregnancy outcomes. This framework motivates the further application of single-cell-resolution approaches to refine models of congenital Chagas pathogenesis and the systematic analysis of maternal peripheral blood during pregnancy to enable early risk stratification and the development of predictive tools for adverse outcomes.

## Introduction

*Trypanosoma cruzi* is the etiological agent of Chagas disease, a parasitic infection responsible for 12,000 deaths annually^1^. The infection is endemic in the Americas and can be acquired through various routes, including vectorial transmission, oral ingestion, blood transfusion, organ transplantation, laboratory accidents, and congenital transmission^2^. Congenital transmission is especially important as it accounts for approximately 22% of new Chagas cases^3,4^, with 5% of infected mothers transmitting the infection to the newborn^5^. Notably, this silent mode of transmission, where *T. cruzi* infection is passed from generation to generation, has become the primary source of new infections in non-endemic regions, presenting a global public health challenge^5–7^.

Congenital transmission of *T. cruzi* poses serious immediate and long-term health risks for infants, and is associated with adverse outcomes in newborns, including low birth weight, prematurity, low Apgar scores, hepatosplenomegaly, anemia, and neonatal death^8,9^. If left untreated, approximately 20-30% of infected infants will, over decades, develop severe and often life-threatening manifestations, such as chronic Chagas cardiomyopathy, megacolon, and/or megaesophagus^10^. Importantly, congenital Chagas disease represents a unique window for therapeutic intervention by monitoring pregnancies of *T. cruzi*-infected mothers during both the prenatal and postnatal periods^11,12^. Despite this opportunity, early intervention remains limited by an incomplete understanding of the molecular mechanisms underlying placental pathology and congenital transmission. Defining these mechanisms is essential not only to understand disease pathogenesis, but also to identify targets for interventions that could prevent placental damage and interrupt transmission. Complementarily, identifying pregnancies at highest risk of adverse outcomes would enable closer monitoring and more effective use of limited clinical resources, rather than simply predicting whether transmission will occur^13,14^.

While the molecular mechanisms of other parasitic congenital infections, such as toxoplasmosis and malaria, are better understood^15,16^, the molecular drivers of congenital Chagas disease remain poorly defined, largely due to limited studies using human samples and a lack of high-throughput, unbiased molecular methods needed to assess placental physiology at the molecular level^17^. To our knowledge, no high-throughput analyses of peripheral blood from naturally *T. cruzi*-infected mothers have been reported, in the context of congenital transmission. Although placental tissue is crucial as the site of congenital transmission, only one study has used high-throughput transcriptomic analysis to examine a limited number of placental tissues from naturally infected pregnancies, and it did not include mothers who transmitted the parasite. Notably, the most recent studies analyzing a larger number of placental tissues from naturally infected mothers were conducted decades ago and were restricted to pathological assessment and low-resolution microscopy^18–20^.

Here, we conducted a comprehensive analysis of placental tissue and peripheral blood from *T. cruzi*-infected mothers who did or did not transmit the parasite to their infants, alongside uninfected controls, to identify molecular drivers of congenital Chagas disease. Combining bulk RNA-seq of blood and placenta, multiplex serum proteomics, and the first spatial transcriptomic map of placental tissue in *T. cruzi* infection, our study provides new insights into the mechanisms of congenital transmission. We highlight the profound impact of *T. cruzi* infection on placental physiology, regardless of whether the parasite successfully crosses the placental barrier.

Particular attention is given to the role of immune response in influencing placental tissue damage, and the potential use of peripheral immune markers to assess the risk of congenital transmission. Our data suggest a paradigm shift from a transmission-centered model to one that recognizes infection-driven placental damage as a spectrum and a primary determinant of adverse pregnancy outcomes. We identified markers predictive of transmission, which might transform the way we monitor and manage Congenital Chagas.

## Results

### *T. cruzi* Infection Broadly Alters Placental Gene Expression

To uncover the impact of maternal *T. cruzi* infection on human placental tissue, we performed bulk RNA sequencing on tissue samples from the placentas of infected mothers (n = 11) and compared them to those from uninfected controls (n = 9). Principal component analysis (PCA) of gene expression variance revealed that maternal *T. cruzi* infection alters placental gene expression dramatically when compared to uninfected mothers, with a significant separation between both groups (PERMANOVA, p= 0.001, R2 = 0.31) **(Fig. 1a)**. Differential gene expression analysis confirmed these dramatic differences, with 3,698 genes downregulated and 3,812 upregulated (fold change ≥ 1.5, FDR ≤ 0.05) **(Fig. 1b, Supplementary File 1)**. Boxplots of the top 20 ranked genes revealed distinct expression patterns, clearly separating infected from uninfected samples **(Fig. 1c)**.

**Figure 1.**
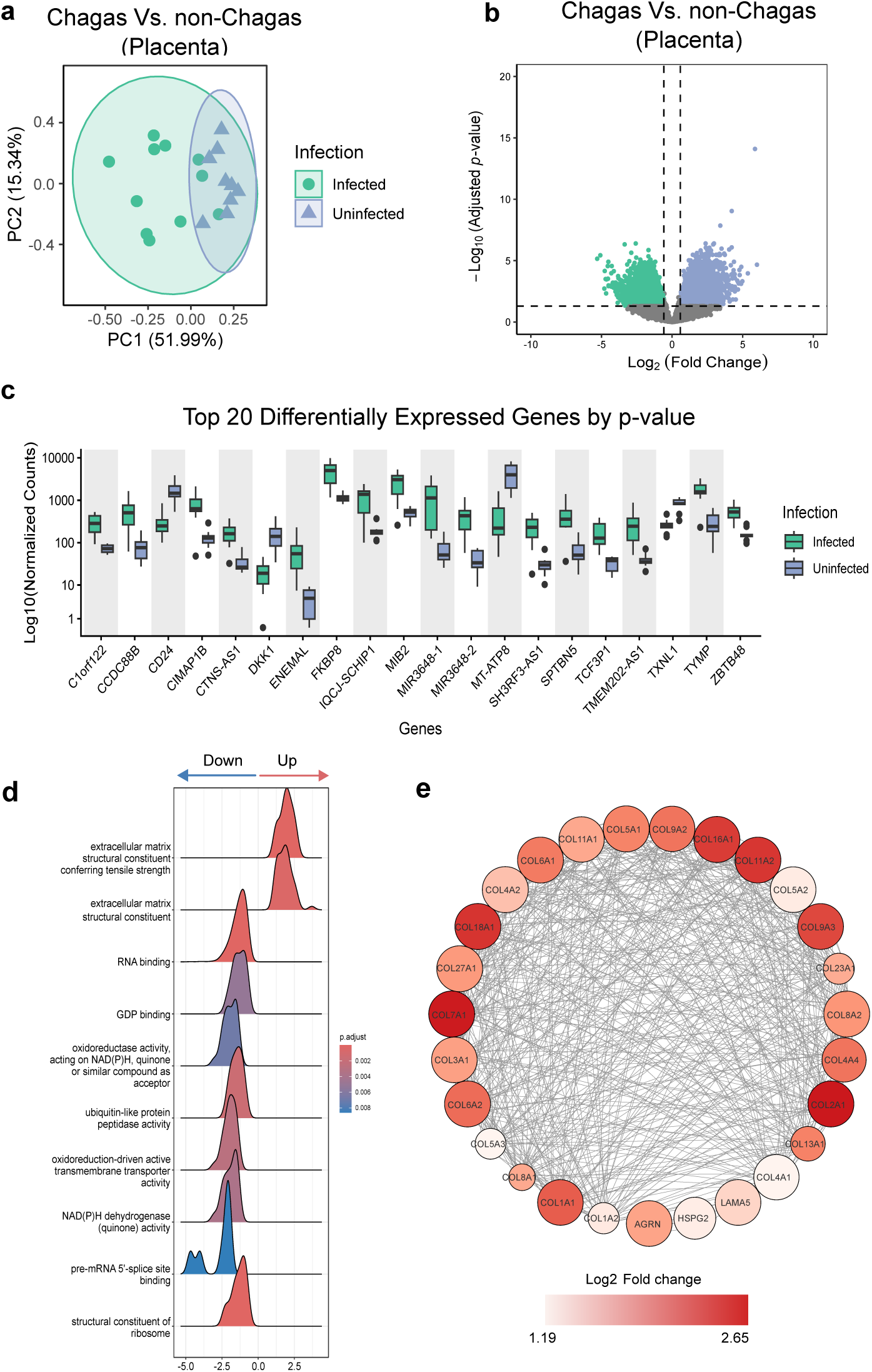
*T. cruzi* infection induces extracellular matrix remodeling in placental tissue, reflecting active local damage. (a) PCA of the top 500 most variable genes across samples. Samples are colored by class (Infected, Uninfected). Ellipses represent one standard deviation from the group mean, assuming a multivariate normal distribution. (b) Volcano plot showing upregulated and downregulated DEGs between placental tissues from infected versus uninfected mothers. (c) Top 20 DEGs ranked by adjusted p-value visualized with normalized counts estimated using the DESeq2 algorithm. (d) Ridge plot illustrating the top 10 enriched molecular functions. Results are based on GSEA performed using the gseGO function from clusterProfiler (version 3.0.4). The blue-to-red gradient represents adjusted p-value distribution corrected by the Benjamini-Hochberg method. Only categories with adjusted p-values (padj) < 0.05 are shown. The X-axis represents the log2 fold change. (e) STRING association network based on manually curated database interactions, summarizes ECM-related components. Node color (white to red gradient) reflects increasing log2 fold change, with larger node size indicating lower FDR, and edges indicate coexpression relationships.

Due to the large number of differentially expressed genes (DEG), we performed Gene Set Enrichment Analysis (GSEA) to evaluate the major functional pathways involved, rather than focusing on individual genes. Besides multiple molecular functions, GSEA identified “extracellular matrix (ECM) structural constituents” as the top-enriched molecular function in placental tissue from *T. cruzi*-infected mothers **(Fig. 1d, Supplementary File 2)**. The increased ECM structural components included the upregulation of 27 transcripts involved in collagen synthesis (**Fig. 1e)**, and six related to elastic fiber assembly (VTN, EMILIN1, FBLN2, ELN, HSPG2, AGRN). The specific gene expression changes suggest that extensive ECM remodeling is associated with infection, potentially reflecting tissue repair in response to damage.

Additionally, *T. cruzi* infection led to broad suppression of key cellular processes, including translation (e.g., structural components of the ribosome, translation initiation and regulatory factors, ribosome binding), energy metabolism (e.g., NADH dehydrogenase activity, oxidoreductase activity, electron transport, and proton/ATP-dependent transporters), and cell cycle regulation (e.g., cyclin-dependent kinase regulators) (**Supplementary File 3**). These transcriptional alterations may indicate a cellular state consistent with hypoxia or stress-induced cell cycle arrest. Together, these findings suggest that maternal *T. cruzi* infection, regardless of whether the parasite is transmitted to the fetus/newborn, profoundly disrupts placental gene expression.

### *T. cruzi* Congenital Transmission is Associated with Distinct Markers of Tissue Damage

While *T. cruzi* infection alters placental gene expression regardless of transmission status, specific expression patterns may distinguish transmitter from non-transmitter mothers. Of the 11 placental tissues evaluated from infected mothers, 7 were from transmitter cases, and 4 were from non-transmitter controls. We analyzed differential gene expression between these groups to identify molecular signatures associated with congenital Chagas transmission. PCA indicated a trend of distinct separation between transmitter and non-transmitter mothers. However, one transmitter (ID:10032) clustered closer to the non-transmitter group, leading to a non-significant overall separation (PERMANOVA, p= 0.28, R2 = 0.11)**(Fig. 2a)**. Differential expression analysis identified 298 significantly altered genes (fold change > 1.5, FDR < 0.05), with the majority (78%) being upregulated in transmitter placentas **(Fig. 2b, Supplementary File 1)**. Boxplots of the top 20 ranked genes showed clear separation, corroborating differences between groups **(Fig. 2c)**.

**Figure 2.**
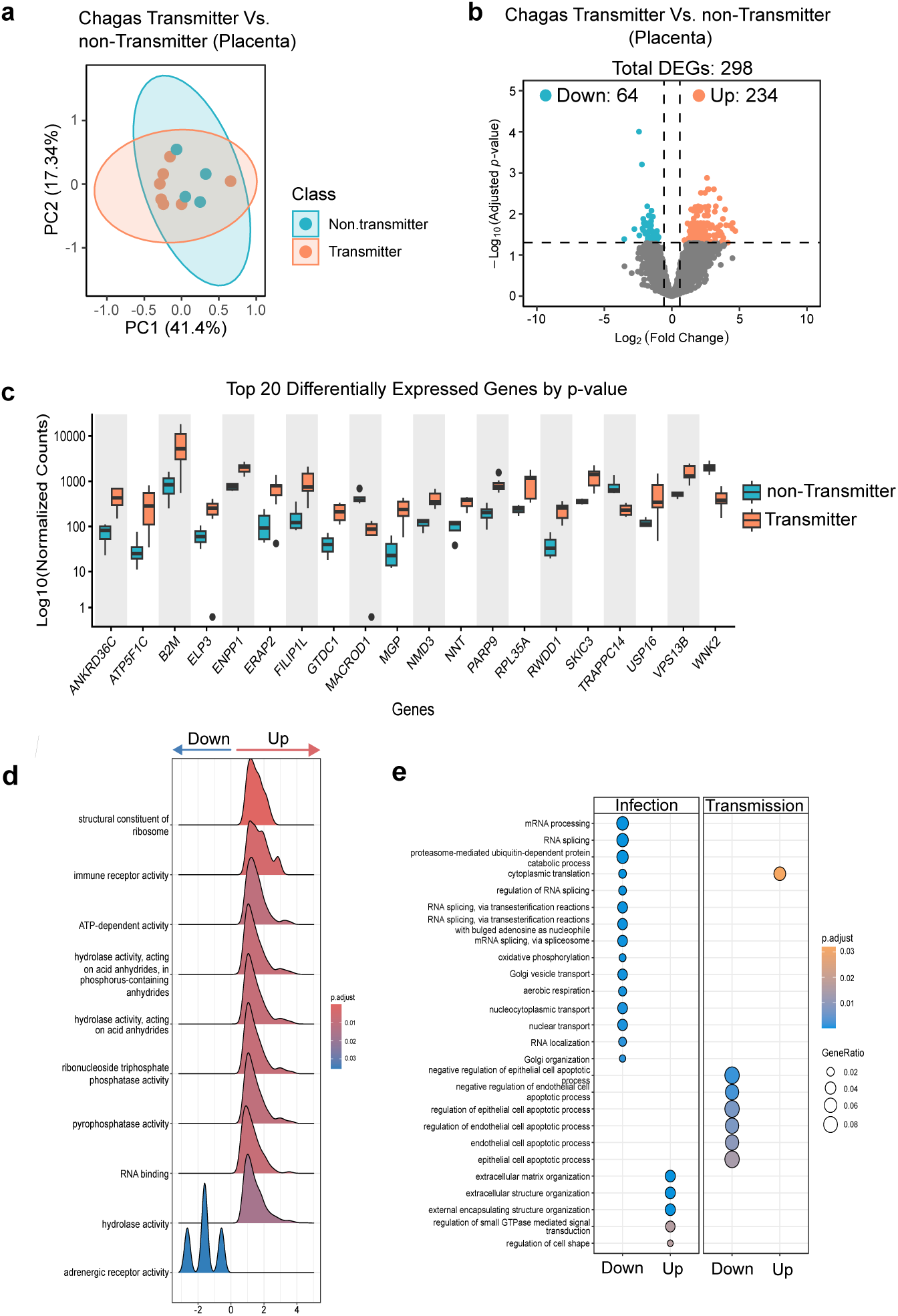
Placental transcriptome reveals exacerbated immune response and apoptosis in mothers transmitting *T. cruzi* to their infants. **(a)** PCA of the top 500 most variable genes across samples. Samples are colored by class (Transmitter, Non-transmitter). Ellipses represent one standard deviation from the group mean, assuming a multivariate normal distribution. **(b)** Volcano plot showing upregulated and downregulated differentially expressed genes (DEGs) in placental tissues from transmitter (orange) vs non-transmitter mothers (blue). **(c)** Top 20 DEGs visualized with normalized counts estimated using the DESeq2 algorithm. **(d)** Ridge plot illustrating the top 10 molecular functions associated with *T. cruzi* transmission. Ridge plots summarize GSEA results obtained using the gseGO function of *clusterProfiler* (version 3.0.4). The blue-to-red gradient represents adjusted p-value distribution corrected by the Benjamini-Hochberg method. The X-axis represents the log2 fold change. **(e)** Enrichment analysis using enrichGO, grouped by fold-change direction (up and down) and comparison group (infection or transmission), performed with the compareCluster function of *clusterProfiler* (version 3.0.4) using DEGs as input. Blue-to-orange gradients represent the adjusted p-value distribution, corrected using the Benjamini-Hochberg method. **(d-e)** Only categories with adjusted p-values (padj) < 0.05 are shown.

GSEA revealed “immune receptor activity” as a key hallmark in transmitter mothers, along with elevated hydrolase and metabolic functions **(Fig. 2d, Supplementary File 2)**. Manual curation identified 47 DEGs potentially linked to tissue injury responses (**Supplementary File 3**). These included elevated expression of immune-inflammatory mediators, such as Toll-like receptors 1 and 7, their downstream effector NF-κB, and a suite of pro-inflammatory genes (e.g., ERAP2, B2M, CTSS, STAT1, MAPK1, FAS, SPP1, IFI44L, NKTR). Disruption of apoptosis and DNA damage repair pathways was also observed, with altered expression of genes implicated in cellular turnover during wound healing (MACROD1, PARP9, CYLD, ATR, SUMO1, FAS, KAT5, DRAM1, CUL5, SOD2). Moreover, upregulation of extracellular matrix components (MGP, FAP, LUM, FILIP1L, PLOD2, FLRT2, DMD) and proliferation-associated genes (MAPK1, FGFR3, ANGPTL1, STAT1, NFKB1, RPS27A, RPL9, RPL21, RPL35A, RPS3A, EIF2A, USP, MED21, CAPZA1, CAND1, TMEM62, DPP4, FOXP116, MAP3K2, COPS4, COPS8) suggest active tissue remodeling and regenerative processes. These molecular signatures point to a dysregulated immune response and increased placental injury in transmitter mothers, which may compromise barrier integrity and facilitate congenital transmission of *T. cruzi*.

To pinpoint transmission-specific alterations, we compared gene ontology pathways overrepresented by the DEGs across two comparisons: infected vs. uninfected, and infected transmitters vs. infected non-transmitters. The analysis revealed distinct patterns between the two comparisons: while maternal *T. cruzi* infection was linked to increased ECM remodeling, congenital transmission was characterized by elevated apoptotic activity. This was evidenced by downregulation of negative apoptotic regulators (e.g., WFS1, MIR126, GATA2, GATA3, and PAK4) **(Fig. 2e)**. Together, these data suggest that the placental tissue of infected transmitter mothers might exhibit a dysregulated syncytiotrophoblast turnover, characterized by altered cell death and insufficient renewal, which might compromises the placental barrier’s integrity, potentially creating gaps that facilitate parasite translocation to the fetus.

### *T. cruzi* infection induces cell and region-specific alterations in the placenta that reveal mechanisms of parasite-induced tissue damage

Given the placenta’s complex architecture and compartmentalized functions, elucidating how *T. cruzi* infection compromises placental barrier integrity requires spatial resolution. We performed a spatial transcriptomic analysis to associate gene expression changes with specific cell populations and native placental histological context. We focused on sections of the chorionic villi, finger-like projections that are essential for nutrient exchange, immune protection, and barrier function, and are predicted to be disrupted by

*T. cruzi* based on ex vivo and in vitro models^21^. Placental tissue was collected from a non-transmitter *T. cruzi*-infected mother and an uninfected control. Samples were matched for maternal age, parity, delivery type, and newborn sex.

First, we compared cell abundance between Chagas and control samples by calculating the proportion of spatial transcriptomic spots for each cell type detected. To identify cell types present in the tissue, we applied Robust Cell Type Decomposition (RCTD) using previously published single-cell RNA-seq data obtained from at-term placental villous tissue (GSE182381) as a reference^22^. The deconvolution analysis resolved 17 cell types, with fetal cytotrophoblasts, endothelial cells, fibroblasts, Fetal Hofbauer Cells (HBCs), and mesenchymal stem cells being the most abundant (**Supplementary Figure 1).** Chagas samples showed higher proportions of HBCs or placental macrophages (1.67-fold increase), proliferative cytotrophoblasts (20.52-fold increase), and syncytiotrophoblasts (17.44-fold increase) and lower proportions of fetal nucleated red blood cells (2.55-fold decrease) and maternal FCGR3A⁺ monocytes (5.23-fold decrease) (**Supplementary Figure 2**). The overgrowth in trophoblasts was also observed through histological analysis, showing reactive trophoblastic hyperplasia.

Notably, HBCs in Chagas samples more frequently colocalized with cytotrophoblasts, fetal endothelial cells, and fetal mesenchymal stem cells (**Supplementary Figure 3**). These findings indicate that the transcriptomic changes previously identified in bulk RNA-seq of placental tissue are likely influenced by altered placental cell composition.

Beyond shifts in cell abundance, transcriptional differences across specific cell types and spatial tissue domains may offer key insights into *T. cruzi*-induced perturbations. To investigate these, we performed a spatial gene expression cluster analysis based on Uniform Manifold Approximation and Projection (UMAP) (**Supplementary Figure 4**).

Of the 10 identified clusters, we focused on two with visually distinct spatial organizations, reflecting divergent tissue microenvironments. The first cluster or spatial region is enriched with trophoblasts, which are known to form a continuous epithelial barrier that makes up the first line of defense against invading pathogens (**Fig. 3a**, green outline). The second spatial region is enriched for mesenchymal stem cells (MSC) overlapping with secondary villi structures, which are part of the villous stroma that represents the second line of defense (**Fig. 3a**, blue outline).

**Figure 3.**
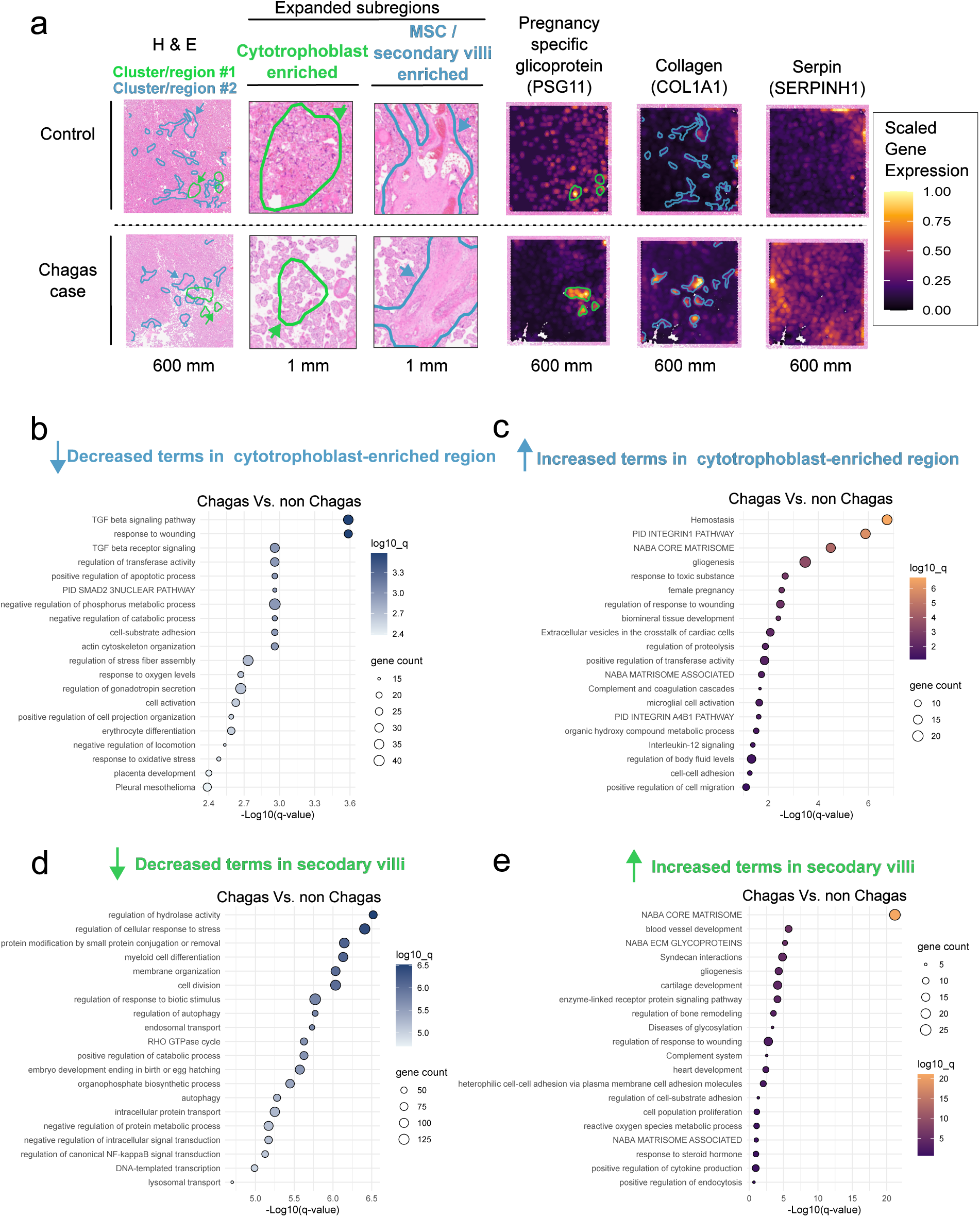
Spatial transcriptomics of placental tissue reveals gene expression changes consistent with bulk RNA-seq findings. **a**, Visium spatial transcriptomics overlaid on its paired H&E-stained section, highlighting tissue subregions and gene expression patterns. Secondary villi enriched in mesenchymal stem cells are outlined in blue; cytotrophoblast-enriched regions are outlined in green. Region annotations were generated using the createImageAnnotations function in SPATA2. Representative genes include *PSG11* (pregnancy-specific glycoprotein), *COL1A1* (collagen), and *SERPINH1*, visualized using the plotSurfaceComparison function in SPATA2. Normalized gene expression is shown on a gradient from low (purple) to high (yellow). **b-e**, Top 20 enriched Gene Ontology terms associated with differentially expressed genes (DEGs) in trophoblast (**b**, decreased; **c**, increased) and secondary villi regions (**d**, decreased; **e**, increased), as identified by Metascape analysis (Supplementary File 4). Bar length reflects gene count per term; color gradient indicates increasing significance as -log₁₀(q-value), adjusted using the Benjamini-Hochberg correction.

By comparing the trophoblast-enriched regions between Chagas (non-transmitter) and non-Chagas placentas, a total of 197 genes were downregulated, and 66 genes upregulated relative to the non-Chagas controls. Downregulated genes were notably associated with tissue repair and wound response pathways, including VASH1, HYAL2, CD9, CLDN1, LYN, VWF, SERPINE2, DUSP, PDGFA, TGFBR1, SMAD3, and CCN2.

Marked repression of both canonical and non-canonical TGF-β signaling pathways was observed, with decreased expression of FN1, SMAD3, TGFBR1, SKI, KLF10, INHBA, CCN2, and ROCK1 (**Fig. 3b)**. Conversely, upregulated genes in the trophoblast-enriched regions included inflammatory markers (SPP1, F13A1, SERPINB2, SRGN, SDC1, SELENOP, SOD1, CLEC3B, JAM3, ANXA6, APOE, IL33, and TRIM25), consistent with placentitis reported in pathology analysis for the Chagas sample. Furthermore, we observed increased potentially anti-tissue damage factors, including several pregnancy-specific glycoproteins (PSG1, PSG3, PSG4, PSG5, PSG6, PSG7, PSG9, PSG11), which play key roles in immune modulation, angiogenesis, and fetal-maternal tolerance, and EBI3 (Epstein-Barr virus induced gene 3), which has anti-inflammatory and immune-modulatory functions (**Fig. 3c)**. These findings highlight that trophoblasts finely tune a balance between tissue repair and pro- and anti-inflammatory responses during infection. This regulatory mechanism may be essential for mitigating *T. cruzi*-induced tissue damage while preserving placental function and integrity.

Within the mesenchymal stem cell-enriched region, placental tissue from *T. cruzi*-infected samples displayed pronounced downregulation of 1,368 genes linked to cell division, transcriptional activity, stress responses, intracellular trafficking, autophagy, and protein homeostasis (**Fig. 3d)**. This transcriptional profile is consistent with a quiescent cellular state, mirroring patterns observed in bulk RNA-seq analyses of infected placental tissue. This hypoactive transcriptomic profile contrasted with the upregulation of only 42 genes, predominantly ECM components, including multiple collagens (*COL1A1, COL1A2, COL3A1, COL4A1, COL5A2, COL6A1, COL6A3, COL11A1,* and *COL14A1*) (**Fig. 3e)**. These results are consistent with bulk RNA-seq findings, which revealed extensive ECM remodeling in response to *T. cruzi* infection. Collectively, the observed shift in MSCs from active proliferative to a matrix-producing, structural phenotype may reflect an adaptive response aimed at preserving the biomechanical integrity of the villous core.

### Peripheral Blood Samples Reflect Physiological Changes in Placental Tissue

To assess whether the localized alterations observed in placental tissue are also reflected systemically in maternal peripheral blood, we analyzed the transcriptome of peripheral blood samples collected right after delivery. This analysis was also performed across two contrasts: infected vs. uninfected, and transmitters vs. non-transmitters.

Unlike in placental tissue, PCA of blood did not reveal clear segregation of samples based on infection or transmission status **(Fig. 4a-b)**. Moreover, differential gene expression analysis identified few, if any, significantly altered transcripts across these comparisons (**Supplementary File 1)**.

**Figure 4.**
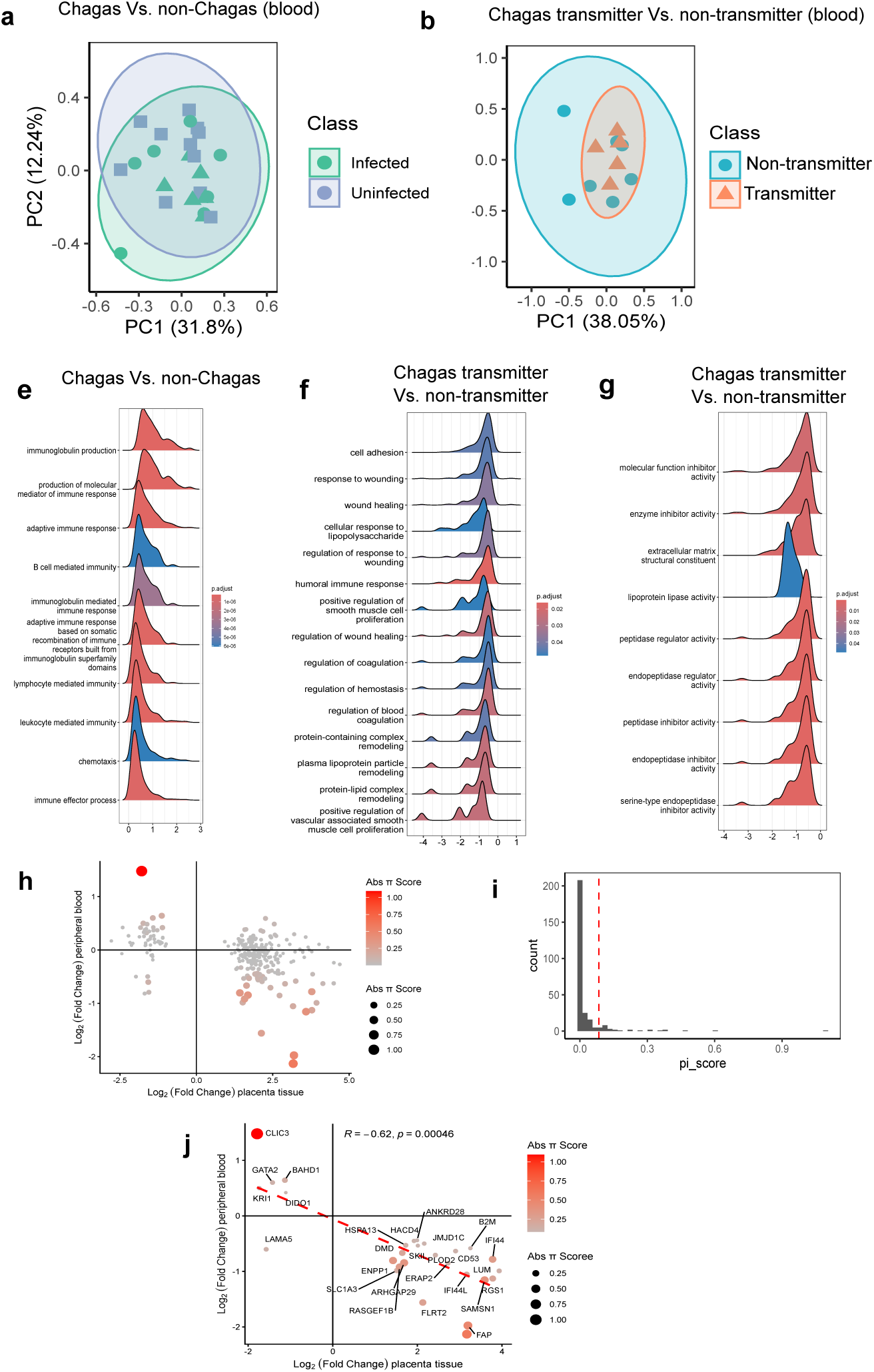
Peripheral blood transcriptome reveals impaired healing pathways in mothers transmitting *T. cruzi* to their infants. **(a)** PCA summarizing gene expression variance between peripheral blood from infected and uninfected mothers. **(b)** PCA summarizing gene expression variance between peripheral blood from transmitter and non-transmitter mothers. **(c)** Ridge plot showing upregulated biological processes linked to *T. cruzi* infection. **(d)** Ridge plot showing downregulated biological processes linked to *T. cruzi* transmission. **(e)** Ridge plot showing downregulated molecular functions linked to *T. cruzi* transmission. **(f)** Association of 288 genes detected as DEG in placental tissue of transmitter mothers and also detectable in the peripheral blood. Both color gradient and size scale represent the absolute π-score. **(g)** Histogram of the distribution of absolute π-score. Vertical lines represent the 90^th^ percentile cutoff. Correlation of the fold change detected in placental tissue (X-axis) and peripheral blood (Y-axis). Linear correlation (red dashed line). Spearman test, Rho: -0.61822, S = 6570, p-value = 0.0004631.

Due to the limited number of significant DEGs, we performed GSEA to uncover more subtle biological alterations. By ranking genes by effect size rather than arbitrary significance thresholds, GSEA identifies enriched pathways that might otherwise be masked by modest transcriptional shifts in peripheral blood. Comparing T. *cruzi*-infected individuals (n = 13) with healthy controls (n = 11), we observed peripheral blood transcriptional signatures indicative of heightened adaptive immune activity, consistent with the expected immune response to parasitic infection **(Fig. 4c, Supplementary File 2).**

In contrast to signatures of infection, perinatal transcriptomic signatures of congenital transmission (Infected transmitters, n = 7 vs. infected non-transmitters, n = 6) were associated with a consistent downregulation of various biological processes and molecular functions. Biological process analysis revealed a coordinated reparative and remodeling state integrating homeostatic pathways (**Fig. 4d**). While the ’regulation of homeostasis’ pathway showed active involvement of coagulation and vascular integrity markers (e.g., F2 F7, F11, F12, PLG, CPB2, TFPI, FGB, FGG, PLAU, PLAUR, PROS1, EDN1, VTN, THBS1, PDGFRA), we observed a parallel suppression of the humoral immune response. Downregulated genes in this category, such as C3, IL6, CXCL8, C5, HLA-DQB1, TREM1, DEFA1, DEFA3, DEFA4, CD46, and MBL2, may reflect a mechanism to mitigate immune overactivation. Furthermore, the reduced expression of critical wound healing modulators (e.g., VEGF, FN1, FGF2, PDGFRA, IGF1, THBS1, ITGB3, MMP2, SOD2, and HIF1A) points towards a potential impairment in tissue repair capacity.

Complementary to this, the analysis based on molecular functions highlights that congenital *T. cruzi* transmission is associated with alterations in ECM-related processes, which are critical for tissue healing. We find significant downregulation of ECM structural components in peripheral blood, including 14 collagens (e.g., COL1A1, COL3A1, COL4A2, COL5A1, COL9A2, COL12A1, COL13A1, COL14A1, COL15A1, COL17A1, COL24A1, COL28A1) and three laminins (LAMA3, LAMA4, LAMA5) **(Fig. 4e)**. This analysis also highlighted gene expression changes suggesting elevated systemic peptidase activity, indicated by the downregulation of multiple peptidase inhibitors, particularly serine protease inhibitors (serpins) from diverse clades (e.g., SERPINA3, SERPINA5, SERPINA6, SERPINA7, SERPINB1, SERPINB5, SERPINC1, SERPIND1, SERPINE2, SERPINE3, SERPINF2, SERPING1, SERPINH1) **(Fig. 4e)**.

Serpins regulate protease activity during wound healing by balancing ECM degradation and deposition, ensuring proper injury resolution. Dysregulated serpins can lead to excessive tissue destruction and sustained inflammatory protease activity. Together, these results indicate that peripheral blood capture transcriptomic alterations linked to wound-healing processes, including protease activity, ECM remodeling, and immune response, implicating pathways also highlighted in placental tissue from Chagas-transmitter mothers.

Because our data indicate that peripheral blood may reflect molecular pathways disrupted in the placenta of transmitting mothers, we investigated whether the expression of some genes in the placenta is mirrored in maternal blood. These shared signals could serve as non-invasive biomarkers to assess placental damage and the risk of congenital transmission. To this end, we intersected the 298 DEGs identified in transmitter placentas with the maternal peripheral blood transcriptome, of which 288 were detectable in circulation. As these genes did not reach conventional significance thresholds in blood, we calculated a π-score for each gene, integrating effect size and statistical significance into a single metric. This analysis revealed a subset of genes with elevated π-scores (aggregated significance), indicated by increased color intensity (**Fig. 4f**). Applying a 90th-percentile cutoff, we identified 27 high-priority candidate genes (**Fig. 4g**). Notably, this subset of genes with higher aggregated significance showed a statistically significant inverse correlation between placental and peripheral blood expression (Spearman’s ρ = −0.62; **Fig. 4h; Supplementary File 5**). These genes are involved in ECM organization (DMD, LAMA5, LUM, PLOD2, FLRT2), response to wounding (GATA2, SLC1A3, JMJD1C), interferon signaling (B2M, IFI44, IFI44L, LAMA5), and multicellular organismal-level homeostasis (B2M, DMD, GATA2, SKIL).

Although samples were obtained from separate individuals, the concordant alterations across tissue compartments reinforce their relevance to congenital Chagas pathophysiology. Together, these findings suggest that peripheral blood markers may serve as surrogate indicators of placental integrity and risk of transmission.

### Serum inflammatory biomarkers inform the risk of congenital *T. cruzi* transmission

Our transcriptomic data suggest that peripheral blood can inform about the risk of congenital transmission; however, proteins often serve as more promising biomarkers due to slower turnover and easier detectability. To assess whether maternal serum markers can predict the risk of congenital transmission of *T. cruzi*, we profiled inflammatory proteins using the standardized commercial panel Olink® Target 96 Inflammation panel (v.3022). This panel was chosen since our transcriptomic data indicate that transmitter mothers exhibit an altered inflammatory immune response.

The panel includes signal transduction, enzymes, growth factors, surface markers, cytokines, and chemokines linked to inflammatory processes. The evaluated serum samples were obtained from a cohort of 81 infected mothers (38 transmitters and 43 non-transmitters).

Using a Hotelling’s T² test with a threshold of two standard deviations, we identified one outlier (sample ID: 6224), which was subsequently excluded from the modeling analysis. Then, the dataset was randomly split into training (80%) and independent test (20%) cohorts. The most informative variables include five proteins directly linked to inflammation (ADA, IL18, CD8A, IL5, CXCL10), and two associated with tissue remodeling (LAP-TGFβ1, MMP-1). A Random Forest classifier trained on the selected proteins yielded an AUC of 0.845 (95% CI: 0.671-0.961; empirical P = 0.008; **Fig. 5a**). Mean accuracy across 100 cross-validation iterations was 0.76 (**Fig. 5b**), significantly outperforming random classification performance.

**Figure 5.**
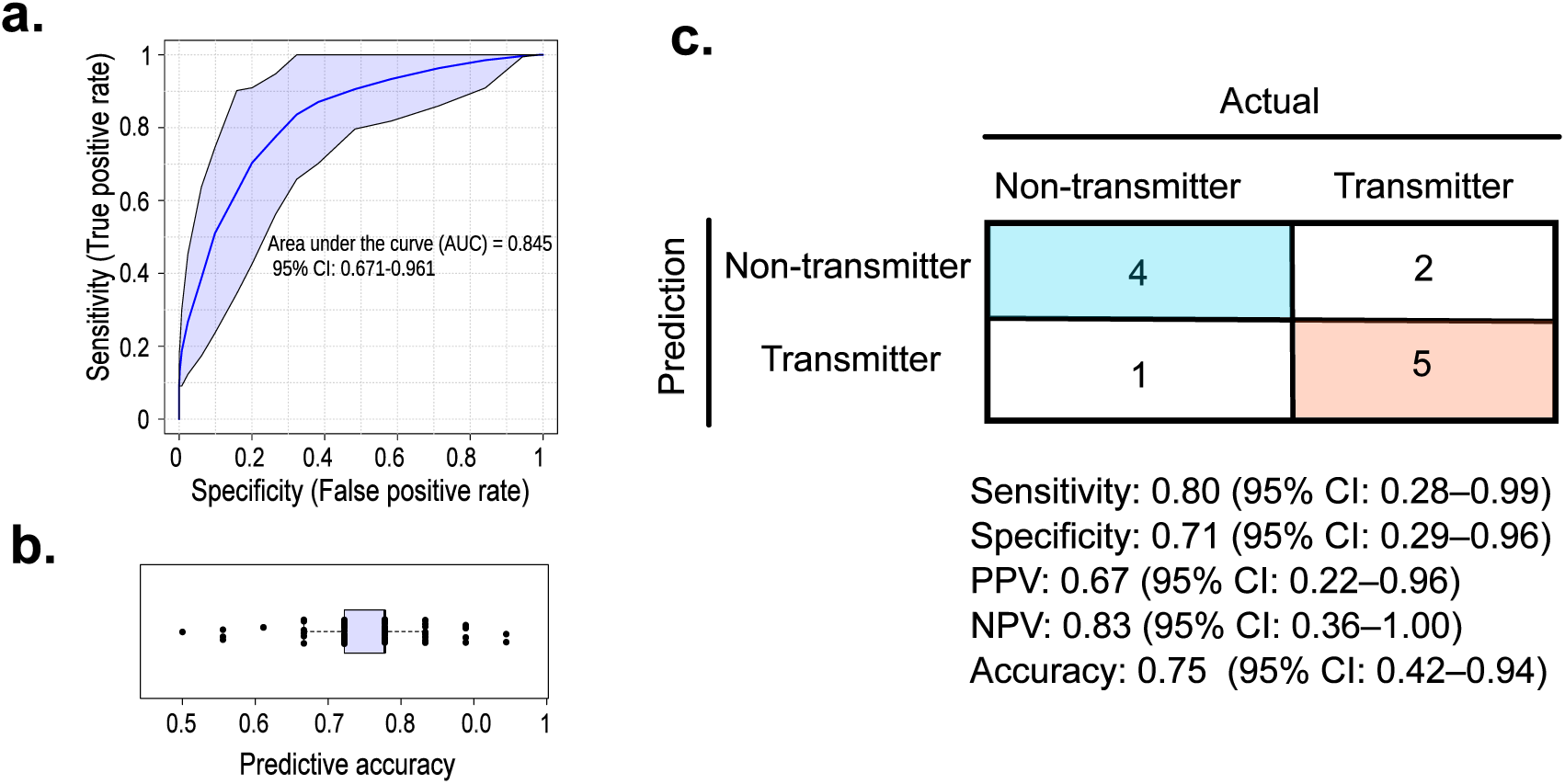
Machine learning model for predicting congenital transmission risk. **a.** Predicted area under the curve (AUC) based on 100 repeated cross-validations, with 95% confidence intervals. **b.** Predicted accuracy from 100 repeated cross-validations, with 95% confidence intervals. **c.** Confusion matrix showing model performance on independent test samples. Positive predictive value (PPV) and negative predictive value (NPV) were estimated using the epiR package, with 95% confidence intervals.

Model evaluation in the independent test cohort demonstrated robust performance: accuracy = 0.75 (95% CI: 0.43-0.95), sensitivity = 0.80 (95% CI: 0.28-0.99), specificity = 0.71 (95% CI: 0.29-0.96), positive predictive value = 0.67 (95% CI: 0.22-0.96), and negative predictive value = 0.83 (95% CI: 0.36-1.00) (**Fig. 5c**). Together, these findings suggest that maternal serum inflammatory profiles capture meaningful variation in congenital transmission risk, highlighting protein biomarkers as a practical tool for clinical stratification.

## Discussion

The lack of high-throughput analyses in naturally infected human samples has limited our understanding of the pathogenesis driving congenital Chagas transmission and hindered improvements in its clinical management. In this study, we employed integrative high-throughput analyses in both peripheral blood and placenta tissue from naturally infected mothers from Bolivia, the country with the highest prevalence of congenital Chagas worldwide. Our findings indicate that maternal *T. cruzi* infection elicits a marked physiological imbalance in the placenta regardless of whether congenital transmission occurs, potentially explaining adverse neonatal outcomes in newborns from non-transmitter mothers^8^. On the other hand, cases of congenital transmission are distinguished by altered inflammatory response and tissue remodeling, suggesting a maladaptive, non-resolving tissue healing response. Remarkably, our findings suggest that peripheral blood profiles offer clinical predictive value to assess the risk of congenital transmission.

Given the placenta’s central role in fetal development and protection, we investigated *T. cruzi*-induced transcriptomic alterations in the local placenta tissue. Our data show widespread transcriptional remodeling involving thousands of genes, in contrast to the limited changes reported previously by Juiz et al. ^23^ This difference may reflect the inherent challenges of working with pooled samples, although other factors such as host ancestry or parasite diversity could also explain this difference. Functional analysis revealed two dominant alterations in placentas from *T. cruzi*-infected mothers: cellular quiescence and extensive ECM remodeling. Consistently, spatial transcriptomics highlighted that both features primarily converge in secondary villi enriched in MSCs, which normally support fetal capillary formation^24^. These alterations indicate a widespread structural remodeling and placental damage in T. cruzi-infected mothers. While such ECM thickening may restrict the parasite from crossing the placental barrier, excessive ECM accumulation likely impairs maternal-fetal exchange^25^, providing a mechanistic framework for recent associations between *T. cruzi* infection and adverse fetal outcomes, such as fetal growth restriction, which appear to be independent of transmission status^8^.

To identify the cellular mediators underlying the gene expression changes observed in the placental tissue of *T. cruzi*-infected mothers, we used spatial transcriptomics-based cell deconvolution. This analysis revealed expansion of HBCs (fetal macrophages that act as inflammatory sensors), consistent with previous in vitro findings^25^. HBCs colocalize with cytotrophoblasts in inflammatory and immunomodulatory niches, indicative of immune-trophoblast crosstalk, potentially driven by both local infection and maternal systemic inflammation. HBCs were also significantly colocalized with MSCs; HBCs typically promote angiogenesis and tissue repair^26^, and their chronic activation may amplify MSC-driven fibrosis and tissue damage^27,28^. Furthermore, we observed a marked increase in proliferative cytotrophoblasts and syncytiotrophoblasts, indicating intensified trophoblast turnover. This suggests a compensatory remodeling response that preserves syncytiotrophoblast integrity, which is known to be altered by *T. cruzi* infection^29,30^. These results indicate that during *T. cruzi* infection, preserving placental integrity requires a balanced and coordinated immune and tissue-repair response.

Although placental protective mechanisms typically prevent congenital transmission in *T. cruzi*-infected mothers, they fail in approximately 5% of pregnancies. Here, we show that *T. cruzi*-infected transmitting mothers are biologically distinct from non-transmitting mothers. While maternal infection induces widespread alterations in the placental transcriptome, congenital transmission is associated with a limited but specific set of changes. Transmission is characterized by an intensified local immune response, including enhanced pattern-recognition and innate immune activation, heightened pro-inflammatory and type I/II interferon signaling, and increased antigen processing and presentation. Our data suggest that congenital Chagas transmission is favored by a heightened inflammatory milieu, similar to congenital toxoplasmosis ^31^ ^32^ and malaria ^33–35^. This immune imbalance is accompanied by dysregulated apoptotic activity, evidenced by the downregulation of negative regulators of apoptosis and increased expression of apoptotic triggers, such as FAS. The excessive or unbalanced cell death might suggest impaired regenerative proliferation during wound healing^36^. Additionally, placentas from *T. cruzi* transmitter mothers exhibited exacerbated activation of ECM pathways, suggesting pathological ECM accumulation. Further histological and protein-based analyses will be required to characterize the potentially excessive ECM deposition in transmitter mothers. In summary, transcriptional signatures consistent with persistent inflammation, dysregulated cell death, and excessive ECM activation indicate that placentas from *T. cruzi* transmitter mothers likely undergo a maladaptive, non-resolving wound-healing response.

The observation that placental tissue distinguishes transmitter from non-transmitter mothers raises the question of whether these local tissue alterations are also reflected at the systemic level in peripheral blood. If so, peripheral blood analyses could have important clinical value for monitoring the risk of congenital Chagas disease. We hypothesize that systemic postpartum physiological changes masked individual gene-level differences, although GSEA effectively uncovered coordinated transcriptional changes. In blood, transmitter mothers exhibited alterations in genes related to wound healing, ECM components, and humoral immune response, consistent with previously discussed pathways altered in the local placental tissue. Markers of vascular integrity were notably increased in the peripheral blood of transmitting mothers, an observation previously reported in *T. cruzi*-infected patients^37,38^. Whether the extent of systemic vascular involvement is associated with placental vascularity and the risk of congenital transmission remains to be clarified. Notably, the effect sizes of DEGs detected in the placenta were inversely correlated with their effect sizes in the peripheral blood of transmitter mothers. Most of these genes participate in ECM organization, response to wounding, interferon signaling, and multicellular organismal-level homeostasis.

Although blood and placental tissue are distinct compartments, they are not completely independent. Circulating immune cells can infiltrate the placenta, and placental cells can be shed into the maternal circulation, for example, during trophoblast turnover. In addition, these compartments can indirectly communicate through cytokines and other signaling molecules, which may partially explain the observed association in gene expression profiles between blood and placental tissues. These findings indicate that peripheral blood might reflect some molecular alterations occurring in the placenta and support its potential use for developing simple, minimally invasive clinical biomarkers to assess the risk of congenital transmission and potentially other adverse outcomes in the context of congenital Chagas.

Our results indicate that tissue remodeling, damage, and inflammatory signatures differ between Chagas-transmitting and non-transmitting mothers, with systemic peripheral blood transcriptomes reflecting alterations observed in the local placental tissue.

Building on this observation, we evaluated whether circulating inflammatory markers could predict the risk of congenital transmission. Our machine learning-based analysis, using a panel of standard inflammatory markers, identified a subset of seven proteins that effectively classify transmitter and non-transmitter mothers (80% sensitivity and 71% specificity). These findings introduce a novel framework for monitoring and managing Chagas disease during pregnancy. Future studies incorporating additional proteins, including new markers of tissue repair and inflammatory modulators described here, may further improve predictive performance. Moreover, longitudinal analyses that include prenatal samples will be essential to identify the best time point to predict adverse outcomes.

This study provides important insight into the potential processes underlying congenital Chagas, but it is subject to some limitations. Although this study represents the first transcriptomic characterization of congenital Chagas disease to include transmitting mothers and to integrate analyses of both peripheral blood and placental tissue, the transcriptomic sample size remains limited. Larger, expanded analyses will be essential to validate our findings. Although most blood and placental samples were obtained from different individuals, the consistency of the molecular signatures across datasets supports the robustness of our conclusions. Future studies incorporating larger cohorts and paired blood-placenta samples from the same patients will be critical to determine whether peripheral blood profiles can reliably serve as surrogates for placental alterations. In addition, we present the first spatial transcriptomic analysis of congenital Chagas disease, offering novel insights into the tissue-level organization of host responses to *T. cruzi*. Integrating spatial approaches with single-cell-resolution technologies will be essential to resolve the cellular and molecular mechanisms of parasite-induced placental damage. While blood is readily accessible and easily sampled, placental tissue remains challenging to study due to its more complex sample processing and marked heterogeneity. Comprehensive histopathological analyses across multiple placental subregions, incorporating inflammatory and tissue repair markers highlighted here, will be necessary to fully capture the complexity of placental pathology and refine models of congenital Chagas pathogenesis.

Overall, this study demonstrates that *T. cruzi* infection profoundly alters the placental environment in which the fetus develops, with the most severe dysregulation occurring in transmitting mothers. These alterations appear to exist along a continuum of adverse effects, of which congenital transmission is likely only one outcome, reflecting more extensive tissue damage. Our findings lay the groundwork for leveraging peripheral blood biomarkers to assess the risk of congenital transmission. The molecular mechanisms described here have the potential to guide innovative strategies to monitor and manage congenital Chagas.

## Methods

### Population of study and specimens

Mothers were recruited from Percy Boland Women’s Hospital in Santa Cruz, Bolivia between the 2016 and 2022. Sample collection was approved by the Institutional Review Boards of the University of North Carolina at Chapel Hill (IRB 19-3014) and Hospital De La Mujer Dr. Percy Boland (Protocol 036).

### Maternal diagnosis of *T. cruzi* infection

Maternal *T. cruzi* infection was determined by serological testing. Mothers were initially screened using either the Chagas Detect™ Plus Rapid Test (InBios, Seattle, WA, USA) or the Chagas STAT-PAK® assay (Chembio Diagnostics, Medford, NY, USA).

Seropositive samples were subsequently confirmed using at least two independent assays, including an in-house IgG TESA-blot, enzyme-linked immunosorbent assay (ELISA; CHAGATEST RECOMBINANTE v3.0, product code 1293254; Wiener Lab, Rosario, Argentina), and hemagglutination inhibition assay (HAI; CHAGATEST HAI Screening A-V; Wiener Lab, Rosario, Argentina), in accordance with standard diagnostic criteria.

### Newborn diagnosis of congenital *T. cruzi* infection

Newborns were assessed for congenital *T. cruzi* infection using a combination of molecular and serological methods. Acute infection was initially evaluated by TaqMan quantitative PCR (qPCR) using heel-prick blood samples (300 µL) collected at birth and at 1, 3 and 9 months of age as previously described^39^. Congenital infection was confirmed by detection of parasite-specific IgM using an in-house TESA-blot between 1 and 7 months of age, and by *T. cruzi-*specific IgG serology after 8 months of age, following the expected clearance of maternal antibodies.

### Peripheral blood samples and RNA isolation for RNAseq

Maternal peripheral blood samples were collected during the postpartum recovery phase, immediately after delivery. Whole blood was mixed with three volumes of 1X DNA/RNA Shield (**Zymo Research, Irvine, CA, USA)**) and homogenized by vortexing. Samples were incubated at 4 °C for at least 12 h prior to storage at −20 °C to prevent clogging during downstream processing. Total RNA was isolated using the Quick-RNA Whole Blood Kit (Zymo Research, R1201) with on-column DNase I treatment, according to the manufacturer’s instructions.

### Placental samples and RNA isolation for RNAseq

Placental tissue were collected during the first hour after delivery. A biopsy of about 50 mg of central placenta (2-5 cm from umbilical cord insertion) was collected from the fetal side, the tissue was trimmed to be less than 0.5 cm in at least one dimension and submerged it in 5 volumes(w/v) of RNA*later* solution. Solution was Incubated overnight at 4°C to allow thorough penetration of the tissue.

Total RNA was extracted using TRIzol™ Reagent (Thermo Fisher, 15596018) following mechanical homogenization (1 mL TRIzol per 50 mg tissue). Briefly, RNA was isolated by chloroform phase separation and isopropanol precipitation, followed by a 75% ethanol wash. RNA pellets were resuspended in RNase-free water and incubated at 60°C to for 10 minutes to ensure complete dissolution. RNA was recovered in 50 µL nuclease-free water and stored at -80 °C.

### RNA purification, quantification, and quality assessment

Following RNA isolation, residual genomic DNA was removed by treatment with TURBO DNase (Thermo Fisher Scientific, AM2238), and RNA was subsequently purified using RNAClean XP magnetic beads (Beckman Coulter, A66514). RNA was eluted in 50 µL of nuclease-free water and stored at −80 °C. RNA concentration was measured by NanoDrop spectrophotometry, while RNA purity and residual DNA contamination (≤10%) were assessed using Qubit RNA HS assay (Thermo Fisher Scientific, #Q32857). RNA integrity was evaluated using the DV200 metric using Agilent RNA 6000 Nano Reagent on Agilent 2100 Bioanalyzer (Agilent Technologies), with samples meeting a threshold of DV200 ≥70% for downstream analyses.

### Bulk RNA sequencing

For peripheral blood-derived RNA, rRNA and globin transcripts were depleted using QIAseq® FastSelect™ rRNA HMR (Qiagen, 334386) and the QIAseq® Globin kit (Qiagen, 335377), respectively. For placental tissue-derived RNA, libraries were prepared using the KAPA Hyper RNA Library Preparation Kit with RiboErase HMR (Roche), following the manufacturer’s protocol. In both cases, non-directional RNA-seq libraries were constructed using Novogene’s proprietary library preparation workflow and sequenced on an Illumina NovaSeq 6000 S4 platform using a 2 × 150 bp paired-end configuration.

### Differential gene expression and gene ontology analysis

Differential gene expression and gene ontology analyses were adapted from Duque *et al.* (2025) ^40^. Gene-level raw read counts were obtained using featureCounts (v2.0.1)^41^. All downstream analyses were performed in R (v4.5.1). To stabilize the variance across expression levels, raw counts were transformed using the variance stabilizing transformation (VST) implemented in DESeq2 and used for principal component analysis (PCA) with the prcomp function in base R. PCA results were visualized using autoplot from ggfortify (v0.4.19)^42^.

To assess differences in multivariate gene expression profiles between groups, permutational multivariate analysis of variance (PERMANOVA) was applied to the VST-based distance matrix using the adonis2 function from the vegan package (v2.7-2)^43^. Statistical significance was evaluated using permutation testing, and a fixed random seed was set to ensure reproducibility.

Differential gene expression analysis was performed using DESeq2 (v1.48.2)^44^, which normalizes read counts for sequencing depth and RNA composition, estimates gene-wise dispersion, and fits a negative binomial generalized linear model. Genes with fewer than three normalized counts per sample were filtered out prior to analysis. Covariates, including maternal age and delivery type, were incorporated into the model and evaluated using likelihood ratio tests. Gene-wise significance was assessed using the Wald test, and p-values were corrected for multiple testing using the Benjamini-Hochberg procedure. Genes with a false discovery rate (FDR) < 0.05 and an absolute log2 fold change (|log2FC|) > 0.585 (corresponding to a ≥50% change) were considered significantly differentially expressed.

Gene ontology (GO) enrichment analyses were conducted using the enrichGO and gseGO functions from clusterProfiler (v4.16.0)^45^, using as background all genes retained after low-expression filtering in the DESeq2 analysis. Over-representation analysis was performed with enrichGO using a one-sided Fisher’s exact test applied to DEGs, whereas gene set enrichment analysis (GSEA) was performed with gseGO on genes ranked by log2 fold change using the fgsea algorithm. GO enrichment and GSEA analyses were performed at both the biological process and molecular function levels.

Pathways with BH-adjusted p-values < 0.05 were considered significantly enriched. GSEA results were visualized using the ridgeplot function from the *enrichplot* package^46^, and semantic similarity among enriched GO terms was computed with pairwise_termsim using the Jaccard similarity index.

### Placental samples for Spatial Transcriptome

Whole placental tissue was immersed in 10% neutral-buffered formalin for 24 h. A section was collected from the central placenta perpendicular to the basal plate, approximately 5 cm from the umbilical cord insertion site, with a thickness of ∼0.5 cm. Tissue was processed and embedded in paraffin following standard protocols to generate formalin-fixed, paraffin-embedded (FFPE) blocks. FFPE blocks were sectioned at 5 µm thickness and stained with haematoxylin and eosin (H&E).

Spatial transcriptomic libraries were generated using the Visium CytAssist Spatial Gene Expression for FFPE kit (10x Genomics), according to the manufacturer’s instructions, and sequenced on an Illumina NovaSeq X platform. Sequencing data were processed using Space Ranger (v3.0.1) and aligned to the human reference genome GRCh38 (Ensembl release 98).

Downstream analyses were performed in R (v4.5.1) using Seurat (v5.4.0)^47^, unless otherwise specified. Gene expression counts were normalized using SCTransform, and the top 30 principal components were used to construct the shared nearest-neighbor graph and generate uniform manifold approximation and projection (UMAP) embeddings. Clustering was performed using the FindClusters function with a resolution of 0.9. Batch effects were corrected using Harmony. Cell-type deconvolution was performed using RCTD (v1.9)^48^ in multi-mode, with a reference single-cell RNA-seq dataset derived from term villous placental tissue from healthy donors^49^. Only high-confidence cell-type assignments were retained for downstream analyses.

Spatial regions of interest were manually annotated using the function createImageAnnotations of SPATA2 package (v3.1)^50^. For differential gene expression analysis, raw (unnormalized) spot-level counts from the *Spatial* assay were used. Spots corresponding to the defined clusters/regions were subsetted, and differential expression between conditions (Chagas disease Vs. control) was assessed using DESeq2, using the Wald test within the negative binomial framework. Genes with a false discovery rate (FDR) < 0.05 and an absolute log2 fold change (|log2FC|) > 0.585 (corresponding to a ≥50% change) were considered significantly differentially expressed. DEGs were submitted to Metascape (https://metascape.org) for functional enrichment analysis using default parameters^51^. Metascape was used to identify enriched biological processes, pathways, and molecular functions across integrated annotation sources, and to group related terms based on semantic similarity.

### Serum-based proteomic analysis

Postpartum peripheral blood serum was collected from 81 *Trypanosoma cruzi*-infected mothers, including 38 classified as congenital transmitters and 43 as non-transmitters. Inflammatory protein levels were quantified using the Olink® Target 96 Inflammation panel (v3022; Olink Proteomics), according to the manufacturer’s instructions. Protein abundance was reported as normalized protein expression (NPX) values.

NPX values were normalized for signal intensity and bridged across assay plates using shared reference samples with the OlinkAnalyze R package (v4.3.2). All downstream analyses were performed in MetaboAnalystR (v4.0), with NPX values scaled using Pareto normalization. One sample (ID 6224) was identified as an outlier based on a Hotelling’s T² test (95% confidence interval) during principal component analysis (PCA) and was excluded from subsequent analyses.

The dataset was randomly partitioned into a training set (80%) and an independent test set (20%), using a fixed random seed (1234) to ensure reproducibility. Mothers who transmitted *T. cruzi* were defined as the target class, while non-transmitters served as the reference class. Feature selection was performed using least absolute shrinkage and selection operator (LASSO) logistic regression. Model stability was assessed across 100 cross-validation iterations, and proteins selected in more than 30% of iterations were retained for downstream modeling. A Random Forest classifier was trained on the selected features using 500 trees (ntree = 500). Model performance was evaluated using receiver operating characteristic (ROC) analysis with 100 repeated cross-validations. Statistical significance of the ROC area under the curve (AUC) was assessed using 1,000 label permutations with a significance threshold of α = 0.05. To assess generalization to unseen data, the final model was applied to the independent test set. Performance metrics, including sensitivity, specificity, accuracy, and Cohen’s kappa coefficient, were calculated with 95% confidence intervals using the epiR R package (v2.0.76).

## Data availability

The deep RNA-Seq and spatial transcriptome datasets supporting the conclusions of this article are available in the Gene Expression Omnibus database from the National Center for Biotechnology under accession number GSE311812. The Olink proteomic dataset will be available in PRIDE Archive (ProteomeXchange). All sample and patient IDs included in this manuscript are deidentified and were not known to anyone outside the research group.

## Code availability

All downstream analyses and figures were generated in R. All scripts used to RNAseq, Visium, and Olink analyses are available at https://github.com/mugnierlab/Gutierrez2025

## Acknowledgements

We thank all members of Dr. Gilman’s and Dr. Mugnier’s laboratories for their valuable discussions and support. This study was funded by the National Institute of Allergy and Infectious Diseases (NIAID) of the National Institutes of Health (NIH) under grant R01AI151295 awarded to N.B. The content is solely the responsibility of the authors and does not necessarily represent the official views of the NIH.

## Ethics declarations

S.A.G.G. and R.G. receive salary support from Moderna, Inc.

## Supplementary information

Supplementary Figures:

**Supplementary Figure 1.** Spatial cell type abundance.

**Supplementary Figure 2.** Cell type abundance ratio.

**Supplementary Figure 3.** Cell-to-cell colocalization based on the Jaccard index**. Supplementary Figure 4.** UMAP plot summarizing gene expression variance.

**Supplementary Files:**

**Supplementary File 1:** Differential Gene Expression Analysis DESeq2 results **Supplementary File 2:** GSEA by molecular function and biological processes **Supplementary File 3:** Manually annotated genes for DEG detected in the placental transcriptome of transmitter mothers.

**Supplementary File 4:** Metascape analysis of cytotrophoblast and secondary villi enriched regions for spatial transcriptome.

**Supplementary File 5:** Correlation of blood and placental gene expression transmitter mothers.

Manual classification of placental DEGs from congenital transmitters into four major biological processes based on bulk RNA-seq.

| Category | Gene | Fold change | Description | Role |
| --- | --- | --- | --- | --- |
| Extracellular matrix composition/interaction | FILIP1L | Up (6.49) | filamin A interacting protein 1 like | Involved in cytoskeleton organization and may regulate cell motility. |
|  | PLOD2 | Up (5.37) | procollagen-lysine,2-oxoglutarate 5-dioxygenase 2 | Plays a role in collagen biosynthesis and cross-linking in the extracellular matrix. |
|  | MGP | Up (8.65) | matrix Gla protein | Regulates calcification in the extracellular matrix. |
|  | FAP | Up (9.15) | fibroblast activation protein alpha | Associated with extracellular matrix remodeling and fibrosis. |
|  | LUM | Up (15.28) | lumican | Affects collagen fibril organization in the extracellular matrix. |
|  | FLRT2 | Up (4.37) | fibronectin leucine rich transmembrane protein 2 | Regulates cell adhesion and interactions with the extracellular matrix. |
|  | DMD | Up (3.12) | Dystrophin | Maintains the structural integrity of muscle fibers by connecting the cytoskeleton to the extracellular matrix. |
|  | LAMA5 | Down (-2.97) | Laminin Subunit Alpha 5 | Integral to basement membrane structure and function, contributing to cell adhesion, migration, and differentiation within the extracellular matrix. |
|  | FGFR3 | Down (-3.29) | Fibroblast Growth Factor Receptor 3 | A key regulator in cell growth, differentiation, and repair through fibroblast growth factor signaling. |
| Cell proliferation modulators | MAPK1 | Up (4.12) | mitogen-activated protein kinase 1 | Plays a critical role in the MAPK/ERK signaling pathway, which is directly involved in cell growth and proliferation. |
|  | FGFR3 | Down (-3.29) | fibroblast growth factor receptor 3 | A fibroblast growth factor receptor involved in regulating cell proliferation, differentiation, and apoptosis. |
|  | ANGPTL1 |  | angiopoietin like 1 | Participate in the formation of blood vessel. |
|  | STAT1 | Up (4.06) | signal transducer and activator of transcription 1 | A transcription factor involved in cellular responses to growth factors and cytokines, affecting proliferation. |
|  | NFKB1 | Up (2.54) | nuclear factor kappa B subunit 1 | Central in the NF-κB signaling pathway, which regulates cell proliferation, survival, and immune response. |
|  | RPS27A, RPL9, RPL21, RPL35A, and RPS3A | RPS27A (3.38), RPL9 (4.75), RPL21 (3.54), RPL35A (4.41), and RPS3A(4.05) |  | Ribosomal proteins essential for protein synthesis, indirectly contributing to cell proliferation. |
|  | EIF2A | Up (6.14) | eukaryotic translation initiation factor 2A | Plays a role in translation initiation, critical for cell growth and proliferation. |
|  | USP16 | Up (5.50) | ubiquitin specific peptidase 16 | Regulates histone H2A deubiquitination, important for cell cycle progression and proliferation. |
|  | MED21 | Up (4.25) | mediator complex subunit 21 | Part of the Mediator complex, which regulates transcription of genes involved in cell proliferation. |
|  | CAPZA1 | Up (9.04) | capping actin protein of muscle Z-line subunit alpha 1 | A capping protein involved in actin filament dynamics, important for cell division and proliferation. |
|  | CAND1 | Up (3.45) | cullin associated and neddylation dissociated 1 | Regulates the ubiquitin-proteasome system, influencing cell cycle and proliferation. |
|  | TMEM62 | **Up (3.6)** | transmembrane protein 62 | Associated with cell cycle and growth processes. |
|  | DPP4 | **Up (4.34)** | dipeptidyl peptidase 4 | Known to regulate proliferation in certain cell types through signaling pathways. |
|  | FOXP1 | **Up (2.72)** | forkhead box P1 | A transcription factor linked to cell cycle regulation and proliferation. |
|  | MAP3K2 | **Up (3.41)** | mitogen-activated protein kinase kinase kinase 2 | Involved in MAP kinase signaling pathways that regulate cell proliferation and survival. |
|  | COPS4, COPS8 | **Up(11.18)**<br>**Up(3.43)** |  | Components of the COP9 signalosome, which influences cell cycle and proliferation |
| Inflammation | NFKB1 | **Up (2.54)** | Nuclear Factor Kappa B Subunit 1 | A master regulator of proinflammatory gene expression. |
|  | ERAP2 | **Up (6.60)** | endoplasmic reticulum aminopeptidase 2 | Involved in antigen processing and presentation, which is crucial for activating immune responses, including inflammation |
|  | B2M | **Up (9.53)** | Beta-2-Microglobulin | Plays a role in the MHC class I antigen presentation pathway, directly influencing immune activation. |
|  | TLR1 | **Up (22.6)** | Toll-Like Receptor 1 | Recognizes pathogen-associated molecular patterns (PAMPs) and activates proinflammatory signaling cascades. |
| | TLR7 | **Up (10.53)** | Toll-Like Receptor 7 | TLR7 is a critical innate immune receptor involved in recognition of parasite RNA, induction of IL-12p40 by dendritic cells, and consequent IFN-γ by T lymphocytes. |
|  | CTSS | **Up (3.63)** | Cathepsin S | A cysteine protease involved in antigen processing and the regulation of inflammatory responses. |
|  | STAT1 | **Up (4.06)** | signal transducer and activator of transcription 1 | A transcription factor that mediates responses to proinflammatory cytokines such as interferons. |
|  | MAPK1 | **Up (4.12)** | Mitogen-Activated Protein Kinase 1 | Part of the MAPK signaling pathway, which is critical for proinflammatory cytokine production. |
|  | FAS | **Up (3.97)** | Fas Cell Surface Death Receptor | Mediates apoptosis and inflammatory signaling in immune cells. |
|  | SPP1 | **Up (5.17)** | secreted phosphoprotein 1 | cytokine that upregulates expression of interferon-gamma and interleukin-12. |
|  | *IFI44L* | **Up (8.97)** | Interferon Induced Protein 44 Like | Predicted to be involved in immune response. |
|  | NKTR | **Up (2.35)** | Natural Killer Cell Triggering Receptor | It is present on the surface of natural killer cells and facilitates their binding to targets. Its expression is regulated by IL2 activation of the cells. |
| Apoptosis and DNA damage | MACROD1 | **Down (4.65)** | mono-ADP ribosylhydrolase 1 | Implicated in the DNA damage response via ADP-ribosylation repair pathways. |
|  | PARP9 | **Up (4.43)** | poly(ADP-ribose) polymerase family member 9 | Involved in DNA repair and apoptosis regulation. |
|  | CYLD | **Up (2.22)** | CYLD lysine 63 deubiquitinase | A tumor suppressor that regulates apoptosis and DNA damage responses. |
|  | ATR | **Up (5.51)** | ATR serine/threonine kinase | Key player in the DNA damage response, activating cell cycle checkpoints and repair. |
|  | SUMO1 | **Up (3.55)** | small ubiquitin like modifier 1 | Regulates apoptosis and DNA repair processes through post-translational modifications. |
|  | FAS | **Up (3.97)** | Fas cell surface death receptor | Critical in apoptosis as a death receptor that initiates programmed cell death. |
|  | KAT5 (TIP60) | **Down (-2.56)** | lysine acetyltransferase 5 | Acetylates proteins involved in the DNA damage response and apoptosis. |
|  | DRAM1 | **Up (2.86)** | DNA damage regulated autophagy modulator 1 | Mediates autophagy and apoptosis, particularly in response to p53 activation. |
|  | CUL5 | **Up (16.17)** | cullin 5 | Located in site of DNA damage. |
|  | SOD2 | **Up (4.74)** | superoxide dismutase 2 | Directly or indirectly in producing DNA damage. |

Note: Forty-seven of the 298 unique DEGs were classified into these four major biological processes.

---

**Supplementary Figure 1.**
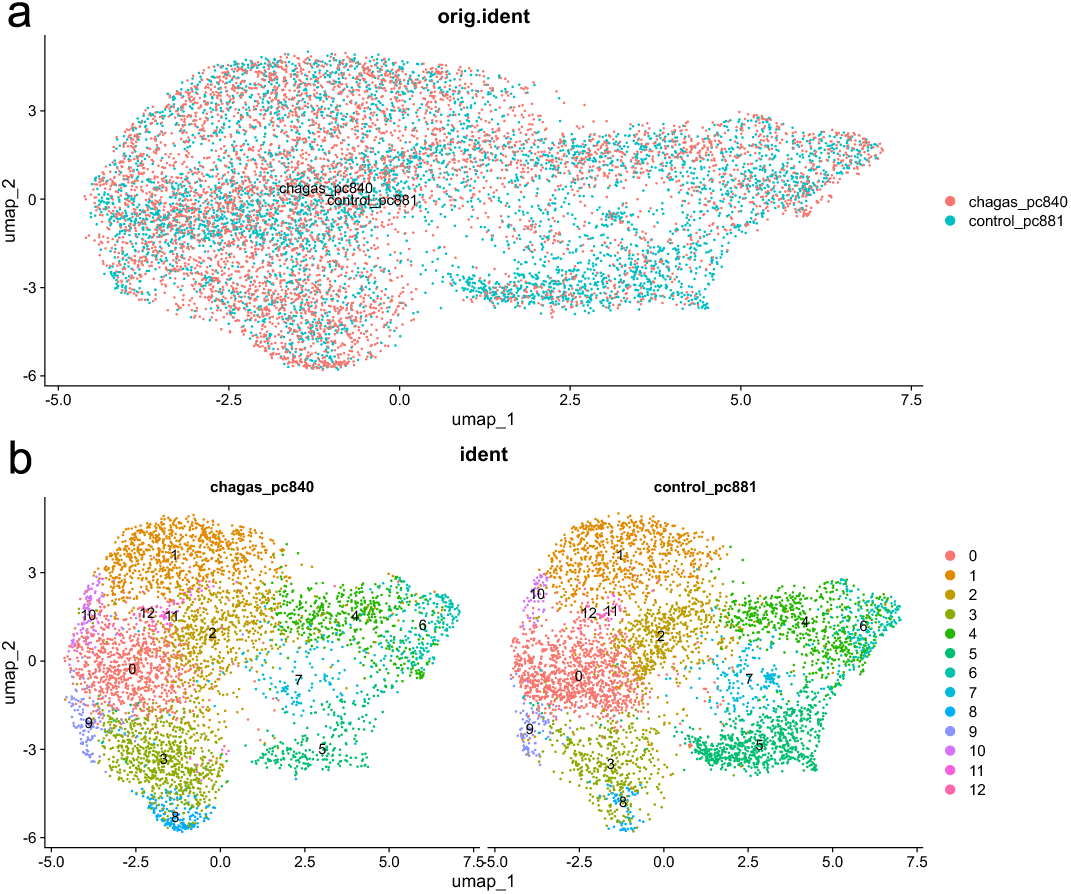
UMAP plot summarizing gene expression variance. **a)** gene variance after sample-to-sample normalization using function FindNeighbors function. **b)** gene expression clusters identified using function FindClusters. Analysis was performed using the Seraut package in R.

**Supplementary Figure 2.**
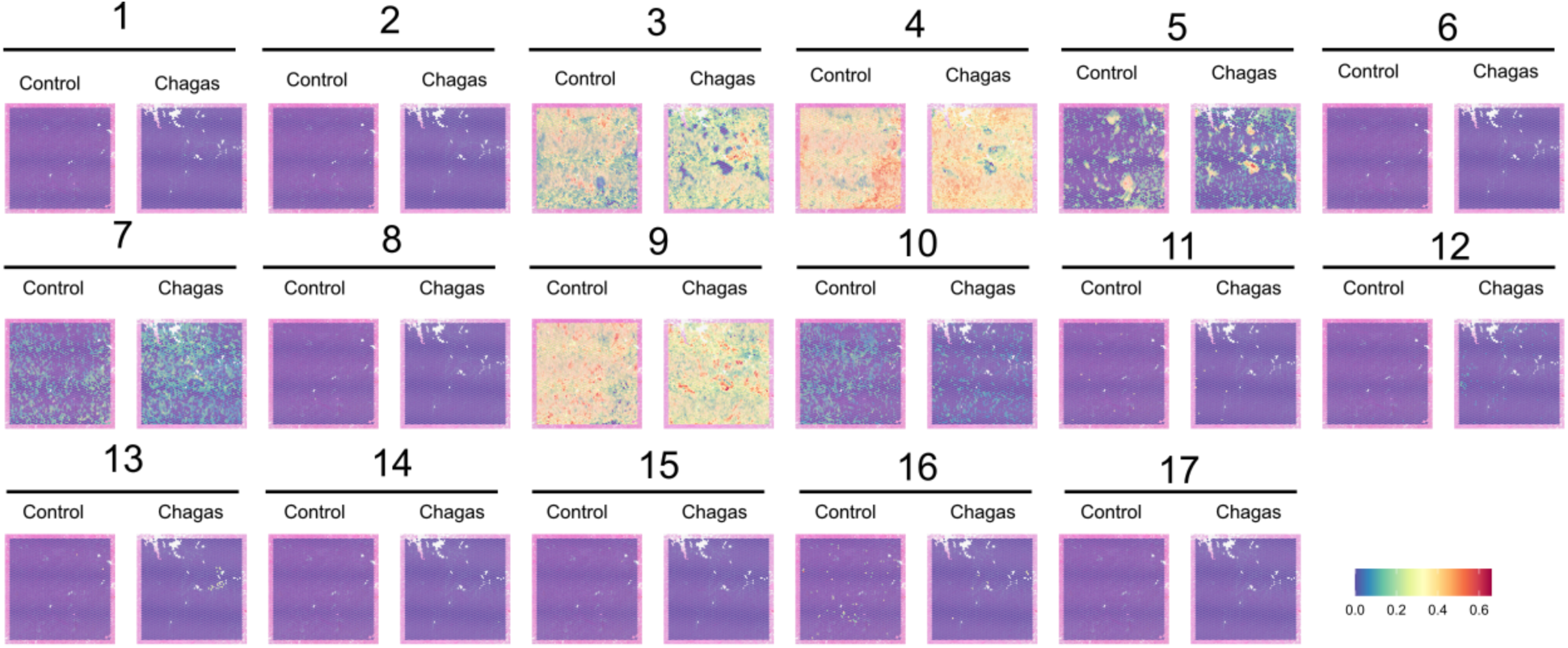
Spatial cell type enrichment per sample. Weights of a cell type in a given spot are represented using a universal scale: low abundance in blue, high abundance in red. Cell types: (1) Fetal CD14+ Monocytes, (2) Fetal CD8+ Cytotoxic T Cells, (3) Fetal Cytotrophoblasts, (4) Fetal Endothelial Cells, (5) Fetal Fibroblasts, (6) Fetal GZMB+ Natural Killer, (7) Fetal Hofbauer Cells, (8) Fetal Memory CD4+ T Cells, (9) Fetal Mesenchymal Stem Cells, (10) Fetal Nucleated Red Blood Cells, (11) Fetal Plasmacytoid Dendritic Cells, (12) Fetal Proliferative Cytotrophoblasts, (13) Fetal Syncytiotrophoblast, (14) Maternal CD14+ Monocytes, (15) Maternal CD8+ Cytotoxic T Cells, (16) Maternal FCGR3A+ Monocytes, (17) Maternal Naive CD4+ T Cells.

**Supplementary Figure 3.**
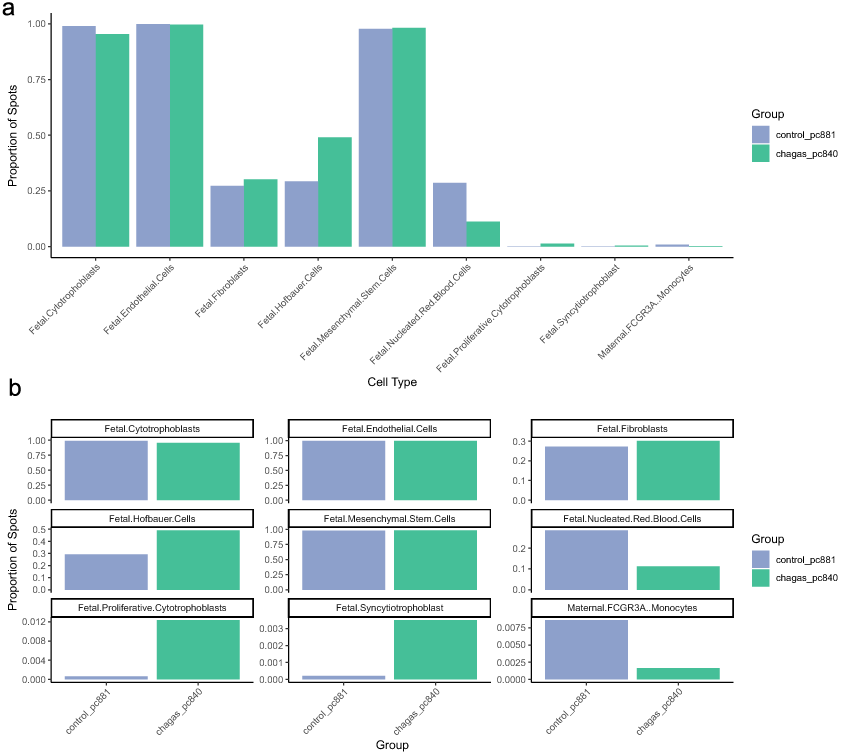
Cell type abundance per sample. A. Bar plot summarizing cell type abundance B. Facet plot summarizing cell type abundance with an independent scale per cell type. Only the nine most abundant cell populations are displayed.

**Supplementary Figure 4.**
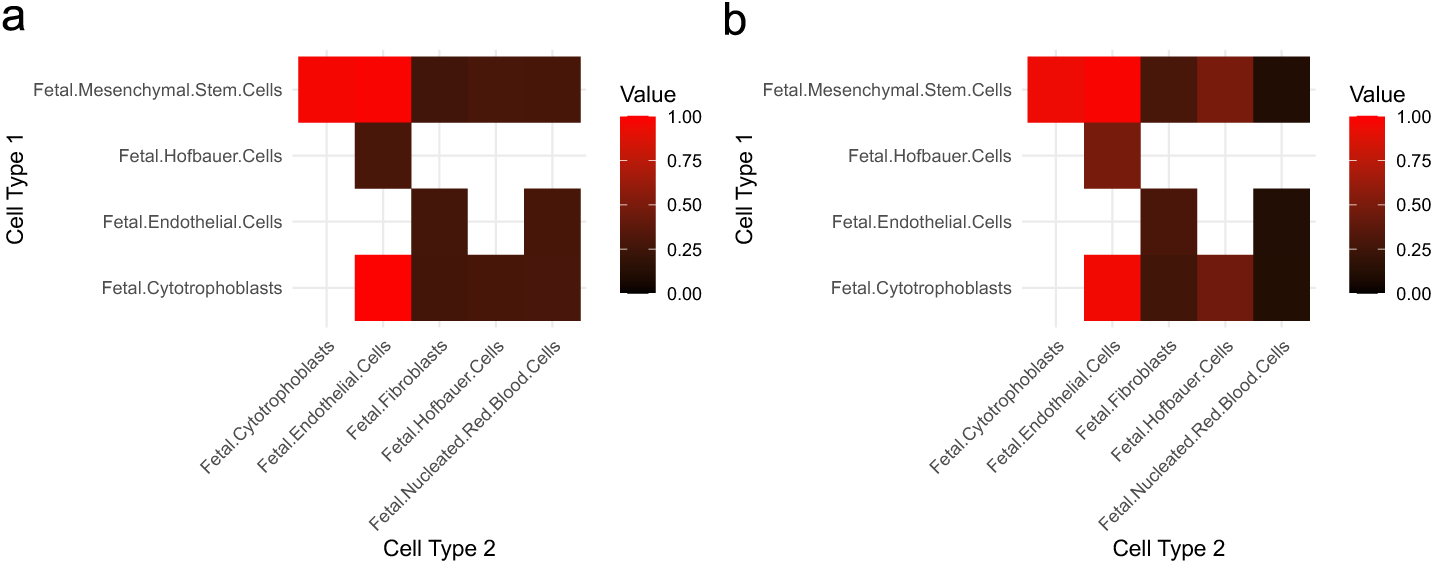
Cell-to-cell colocalization based on Jaccard index**. A.** placenta tissue from uninfected control. **B.** placenta tissue from Chagas patient. Only cells exhibiting at least one colocalization with another cell are displayed.

## References

1. CDC. Parasites - American Trypanosomiasis (also known as Chagas Disease). (2021).

2. WHO. Chagas disease (also known as American trypanosomiasis). https://www.who.int/news-room/fact-sheets/detail/chagas-disease-(american-trypanosomiasis)#:∼:text=Chagas%20disease%2C%20also%20known%20as,cruzi. (2023).

3. 3. WHO. Weekly epidemiological record Relevé épidémiologique hebdomadaire. http://www.who.int/wer (2015).

4. Edwards, M. S. & Montgomery, S. P. Congenital Chagas disease: progress toward implementation of pregnancy-based screening. Current Opinion in Infectious Diseases 34, 538–545 (2021).

5. Howard, E., Xiong, X., Carlier, Y., Sosa-Estani, S. & Buekens, P. Frequency of the congenital transmission of *T rypanosoma cruzi* : a systematic review and meta-analysis. BJOG 121, 22–33 (2014).

6. Avaria, A., Ventura-Garcia, L., Sanmartino, M. & Van Der Laat, C. Population movements, borders, and Chagas disease. Mem. Inst. Oswaldo Cruz 117, e210151 (2022).

7. Messenger, L. A. & Bern, C. Congenital Chagas disease: current diagnostics, limitations and future perspectives. Current Opinion in Infectious Diseases 31, 415–421 (2018).

8. Paternina-Caicedo, A. et al. Trypanosoma cruzi Infection in Pregnancies without Congenital Transmission Is Associated with Reduced Fetal Growth: A Cross-Sectional Study in Argentina, Honduras, and Mexico. The American Journal of Tropical Medicine and Hygiene 111, 64–72 (2024).

9. CDC. Clinical Considerations for Congenital Chagas Disease.

10. Prata, A. Clinical and epidemiological aspects of Chagas disease. The Lancet Infectious Diseases 1, 92–100 (2001).

11. Jackson, Y., Wyssa, B. & Chappuis, F. Tolerance to nifurtimox and benznidazole in adult patients with chronic Chagas’ disease. Journal of Antimicrobial Chemotherapy 75, 690–696 (2020).

12. Morillo, C. A. et al. Randomized Trial of Benznidazole for Chronic Chagas’ Cardiomyopathy. N Engl J Med 373, 1295–1306 (2015).

13. Ribeiro, I. et al. New, Improved Treatments for Chagas Disease: From the R&D Pipeline to the Patients. PLoS Negl Trop Dis 3, e484 (2009).

14. Pecoul, B. et al. The BENEFIT Trial: Where Do We Go from Here? PLoS Negl Trop Dis 10, e0004343 (2016).

15. Pfaff, A. W. et al. Cellular and molecular physiopathology of congenital toxoplasmosis: The dual role of IFN-γ. Parasitology 134, 1895–1902 (2007).

16. Doritchamou, J. et al. Dynamics in the Cytoadherence Phenotypes of Plasmodium falciparum Infected Erythrocytes Isolated during Pregnancy. PLoS ONE 9, e98577 (2014).

17. Rapp, E. & Gold, M. Knowledge Production on Congenital Chagas Disease across Time, Borders and Disciplines: A Comprehensive Scoping Review. TropicalMed 8, 422 (2023).

18. Bittencourt, A. L. Congenital Chagas Disease. Arch Pediatr Adolesc Med 130, 97 (1976).

19. Fernandez-Aguilar, S. et al. [Placental lesions in human Trypanosoma cruzi infection]. Rev Soc Bras Med Trop 38 **Suppl 2**, 84–86 (2005).

20. Lopes, E. R., et al. DOENÇA DE CHAGAS E GRAVIDEZ-Estudo de 50 placentas de gestantes chagásicas crônicas. Rev. lnst. Med. trop. São Paulo (1967).

21. Mayhew, T. M. Turnover of human villous trophoblast in normal pregnancy: What do we know and what do we need to know? Placenta 35, 229–240 (2014).

22. Wang, M. et al. Single-nucleus multi-omic profiling of human placental syncytiotrophoblasts identifies cellular trajectories during pregnancy. Nat Genet 56, 294–305 (2024).

23. Juiz, N. A. et al. Alterations in Placental Gene Expression of Pregnant Women with Chronic Chagas Disease. The American Journal of Pathology 188, 1345–1353 (2018).

24. Zhang, Y., Zhong, Y., Zou, L. & Liu, X. Significance of Placental Mesenchymal Stem Cell in Placenta Development and Implications for Preeclampsia. Front. Pharmacol. 13, 896531 (2022).

25. Bane, A. L. & Gillan, J. E. Massive perivillous fibrinoid causing recurrent placental failure. BJOG 110, 292–295 (2003).

26. Loegl, J. et al. Hofbauer cells of M2a, M2b and M2c polarization may regulate feto-placental angiogenesis. Reproduction 152, 447–455 (2016).

27. Mercnik, M. H., Schliefsteiner, C., Fluhr, H. & Wadsack, C. Placental macrophages present distinct polarization pattern and effector functions depending on clinical onset of preeclampsia. Front. Immunol. 13, 1095879 (2023).

28. Erlandsson, L. et al. Urban air pollution disrupts placental microarchitecture and shifts hofbauer cells towards a pro-inflammatory state. Journal of Environmental Sciences 160, 124–134 (2026).

29. Barbosa, C. G. et al. Congenital transmission of Mexican strains of *Trypanosoma cruzi* TcIa: interaction between parasite and human placental explants. Parasitology 149, 418–426 (2022).

30. Mezzano, L. et al. Chagas disease affects the human placental barrier’s turnover dynamics during pregnancy. Mem. Inst. Oswaldo Cruz 117, e210304 (2022).

31. Gómez-Chávez, F. et al. A Proinflammatory Immune Response Might Determine Toxoplasma gondii Vertical Transmission and Severity of Clinical Features in Congenitally Infected Newborns. Front. Immunol. 11, 390 (2020).

32. Gómez-Chávez, F. et al. Maternal Immune Response During Pregnancy and Vertical Transmission in Human Toxoplasmosis. Front. Immunol. 10, 285 (2019).

33. Suguitan, Jr., A. L., et al. Changes in the Levels of Chemokines and Cytokines in the Placentas of Women with *Plasmodium falciparum* Malaria. J INFECT DIS 188, 1074–7082 (2003).

34. Fried, M. et al. Systemic Inflammatory Response to Malaria During Pregnancy Is Associated With Pregnancy Loss and Preterm Delivery. Clinical Infectious Diseases 65, 1729–1735 (2017).

35. Dobaño, C. et al. High production of pro-inflammatory cytokines by maternal blood mononuclear cells is associated with reduced maternal malaria but increased cord blood infection. Malar J 17, 177 (2018).

36. Landén, N. X., Li, D. & Ståhle, M. Transition from inflammation to proliferation: a critical step during wound healing. Cell. Mol. Life Sci. 73, 3861–3885 (2016).

37. Hassan, G. S., et al. *Trypanosoma cruzi* Infection Induces Proliferation of Vascular Smooth Muscle Cells. Infect Immun 74, 152–159 (2006).

38. Volpini, X., et al. *Trypanosoma cruzi* Infection Promotes Vascular Remodeling and Coexpression of α-Smooth Muscle Actin and Macrophage Markers in Cells of the Aorta. ACS Infect. Dis. 8, 2271–2290 (2022).

39. Gutierrez Guarnizo, S. A., et al. A specific, stable, and accessible LAMP assay targeting the HSP70 gene of *Trypanosoma cruzi*. Microbiol Spectr 13, e00172–25 (2025).

40. Duque, C. et al. Immunologic changes in the peripheral blood transcriptome of individuals with early-stage chronic Chagas cardiomyopathy: a cross-sectional study. Lancet Reg Health Am 45, 101090 (2025).

41. Liao, Y., Smyth, G. K. & Shi, W. featureCounts: an efficient general purpose program for assigning sequence reads to genomic features. Bioinformatics 30, 923–930 (2014).

42. Horikoshi, M. & Tang, Y. Ggfortify: Data Visualization Tools for Statistical Analysis Results. (2016).

43. Oksanen, J., et al. vegan: Community Ecology Package. 2.7-2 10.32614/CRAN.package.vegan (2001).

44. Love, M. I., Huber, W. & Anders, S. Moderated estimation of fold change and dispersion for RNA-seq data with DESeq2. Genome Biol 15, 550 (2014).

45. Guangchuang Yu [Aut, C. clusterProfiler. Bioconductor 10.18129/B9.BIOC.CLUSTERPROFILER (2017).

46. Guangchuang Yu. enrichplot. Bioconductor 10.18129/B9.BIOC.ENRICHPLOT (2018).

47. Satija, R. Seurat: Tools for Single Cell Genomics. 5.4.0 10.32614/CRAN.package.Seurat (2017).

48. Cable, D. M. et al. Robust decomposition of cell type mixtures in spatial transcriptomics. Nat Biotechnol 40, 517–526 (2022).

49. Campbell, K. A. et al. Placental cell type deconvolution reveals that cell proportions drive preeclampsia gene expression differences. Commun Biol 6, 264 (2023).

50. Kueckelhaus, J. et al. Inferring histology-associated gene expression gradients in spatial transcriptomic studies. Nat Commun 15, 7280 (2024).

51. Zhou, Y. et al. Metascape provides a biologist-oriented resource for the analysis of systems-level datasets. Nat Commun 10, 1523 (2019).

